# *In Vivo* Directed Evolution of an Ultra-Fast Rubisco from a Semi-Anaerobic Environment Imparts Oxygen Resistance

**DOI:** 10.1101/2025.02.17.638297

**Authors:** Julie L. McDonald, Nathan P. Shapiro, Amanuella A. Mengiste, Sarah Kaines, Spencer M. Whitney, Robert H. Wilson, Matthew D. Shoulders

**Author notes:** **Author Contributions:** J.L.M., R.H.W., and M.D.S. conceived this work. All authors designed experiments and analyzed data. J.L.M., N.P.S., R.H.W., A.A.M., S.K., and S.M.W. performed experiments. J.L.M., S.M.W., R.H.W, and M.D.S wrote the manuscript. All authors edited the manuscript.

## Abstract

Carbon dioxide (CO_2_) assimilation by the enzyme Ribulose-1,5-bisphosphate Carboxylase/Oxygenase (Rubisco) underpins biomass accumulation in photosynthetic bacteria and eukaryotes. Despite its pivotal role, Rubisco has a slow carboxylation rate 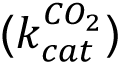 and is competitively inhibited by oxygen (O_2_). These traits impose limitations on photosynthetic efficiency, making Rubisco a compelling target for improvement. Interest in Form II Rubisco from *Gallionellaceae* bacteria, which comprise a dimer or hexamer of large subunits, arises from their nearly 5-fold higher 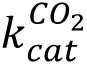 than the average Rubisco enzyme. As well as having a fast 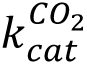 (25.8 s**^−^**^1^ at 25 °C), we show that *Gallionellaceae* Rubisco (GWS1B) is extremely sensitive to O_2_ inhibition, consistent with its evolution under semi-anaerobic environments. We therefore used a novel *in vivo* mutagenesis-mediated screening pipeline to evolve GWS1B over six rounds under oxygenic selection, identifying three catalytic point mutants with improved ambient carboxylation efficiency; Thr-29-Ala (T29A), Glu-40-Lys (E40K) and Arg-337-Cys (R337C). Full kinetic characterization showed that each substitution enhanced the CO_2_ affinity of GWS1B under oxygenic conditions by subduing oxygen affinity, leading to 25% (E40K), 11% (T29A) and 8% (R337C) enhancements in carboxylation efficiency under ambient O_2_ at 25 °C. By contrast, under the near anaerobic natural environment of *Gallionellaceae*, the carboxylation efficiency of each mutant was impaired ∼16%. These findings demonstrate the efficacy of artificial directed evolution to access novel regions of catalytic space in Rubisco.

**Significance:** Given Rubisco’s crucial role in carbon dioxide assimilation, addressing its slow carboxylation rate and oxygen inhibition is a significant challenge. Utilizing one of the fastest known, yet also highly oxygen-sensitive, Rubisco – from the bacteria *Gallionellaceae* – we applied a novel *in vivo* directed evolution pipeline in *Escherichia coli* to discover mutations that specifically enhance carboxylation efficiency under ambient oxygen, a condition distinct from *Gallionellaceae’s* natural semi-anaerobic environment. Our findings underscore the potential of directed evolution to unlock new catalytic capabilities for Rubisco, with implications for both fundamental research and practical agricultural applications.

## Introduction

The enzyme Ribulose-1,5-bisphosphate Carboxylase/Oxygenase (Rubisco) is essential for life on Earth, fixing atmospheric carbon dioxide (CO_2_) to organic carbon during photosynthesis (1). The Rubisco enzyme family has diverse isoforms across phylogeny. Plants, algae, and many mixotrophic and autotrophic bacteria utilize Form I Rubisco, where the enzyme is composed of a large (RbcL) and small (RbcS) subunit as an RbcL_8_RbcS_8_ hexadecamer. Form II and Form III Rubisco, found in bacteria and archaea, lack RbcS and present as arrangements of RbcL dimers (RbcL_2_–RbcL_10_) (2–4). Form II/III Rubisco display lower substrate specificity for CO_2_ over oxygen (O_2_; *S*_C/O_) than Form I enzymes, a deficiency attributed to their lack of RbcS (5).

Despite their widespread prevalence in Nature and essentiality for carbon fixation, Rubisco enzymes display striking inefficiencies. The rate of Rubisco carboxylation is most often in the range of 1–10 reactions per second (6) – much slower than most other enzymes involved in carbon metabolism. The enzyme also reacts promiscuously with O_2_, producing 2-phosphoglycolate (2-PG) that must be recycled via energy-consuming mechanisms, including photorespiration in plants (7) and the 2-PG salvage pathway in bacteria (8). Rubisco’s weak specificity for CO_2_ may be related to the evolutionary emergence of its complex, 5-step, catalytic mechanism in an anoxic atmosphere, prior to the rise of oxygenic photosynthesis (9, 10). The emergence of Earth’s current O_2_-rich atmosphere has necessitated Rubisco to gain selectivity for CO_2_ over O_2_, albeit at a profoundly slow pace (11). Kinetic trends across the Rubisco family suggest that improvements in *S*_C/O_ come at the expense of 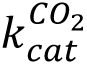 (6, 12, 13), in particular for Form II Rubisco whose particularly high 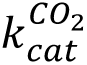 values (>10 s**^−^**^1^) and low *S*_C/O_ (<20 mol/mol) (2) are associated with environments high in dissolved CO_2_ and/or low in O_2_, such as acidic iron-rich soils (14) and the deep sea (15).

A large body of both theoretical and experimental research indicates that improving the efficiency of Rubisco carboxylation can enhance plant photosynthesis (1, 16–20), making Rubisco itself a promising engineering target to enhance crop productivity and yields.

Photosynthetic modeling in crops like rice, wheat and soybean shows such improvements require combined improvements in the 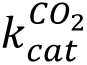, CO_2_-affinity 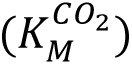 and *S*_C/O_ of Rubisco within the aerobic chloroplast environment (16, 19). While several directed evolution studies on non-plant Rubisco have identified mutations that improve 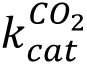 (21–26), anaerobic 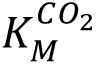 (21, 25, 27–31), and *S*_C/O_ (21–23) of Rubisco, in most instances they overlook how the enzyme’s sensitivity to O_2_ inhibition is impacted (21, 24–30, 32–34). This knowledge is critical to ascertaining the biologically-relevant aerobic 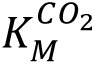(21% O_2_) and carboxylation efficiency 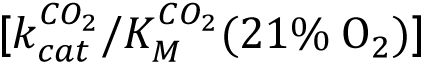 values needed to simulate Rubisco performance in an oxygenic photosynthesis environment (17, 19).

In addition to Rubisco engineering efforts, several studies have sought to profile the natural catalytic landscape of the Rubisco family (2, 35, 36). Form II Rubisco enzymes from the semi-anaerobic bacterial family *Gallionellaceae* have garnered attention from such profiling efforts and from metagenomic analyses (2, 37), as they display some of the fastest Rubisco-mediated carboxylation rates ever measured (15–29 s**^−^**^1^) (2, 38, 39). Interestingly, *Gallionellaceae* Rubisco, while very fast, also undesirably have very high affinities for O_2_ (38). The combination of these extreme and unique properties (fast carboxylation kinetics and high sensitivity to O_2_) position *Gallionellaceae* Rubisco as a novel starting point for directed evolution, with the potential to uncover distinct loci important for mitigating the enzyme’s reactivity with O_2_.

Here, we describe an *in vivo* laboratory directed evolution pipeline for Rubisco that leverages the targeted mutagenesis tool MutaT7 (40–47). This method sidesteps the low-throughput, shallow, and labor-intensive *in vitro* mutagenesis approaches employed in prior Rubisco directed evolution campaigns (21–29, 31–34, 48). MutaT7 is a targeted, genome-encoded mutagen composed of a deaminase fused to a T7 RNA polymerase (T7 RNAP). MutaT7 processively and selectively mutates any DNA region that is flanked by a T7 promoter and terminator. The high mutagenesis frequency (40–47) of MutaT7 positions it as a useful tool for deep sampling of the Rubisco sequence landscape. We apply MutaT7 (42) in tandem with Rubisco-dependent *Escherichia coli* (RDE/RDE2) (23) to evolve a hexameric *Gallionellaceae* Rubisco (termed GWS1B, PDBID: 5C2G) originating from a groundwater sample (37). Our campaign converged on multiple “winner” mutants encoding amino acid substitutions positioned near the active site and oligomeric interfaces. Consistent with the aerobic selection pressure we applied, kinetic analysis revealed that some of the mutations imparted O_2_ resistance on GWS1B variants that also retained their other beneficial catalytic features, thereby enhancing aerobic carboxylation efficiencies.

In sum, by beginning our evolution campaign with an exceptionally fast but also remarkably O_2_-sensitive natural Rubisco, we were able to uncover Rubisco variants that exist in a region of catalytic space not previously observed for Rubisco. These results strongly motivate continued Rubisco engineering efforts that can improve the enzyme beyond its natural capabilities.

## Results

### Biochemical characterization of GWS1B Rubisco

Kinetic characterization of both wild-type and C-terminal polyhistidine-tagged GWS1B showed that the epitope tag did not impact 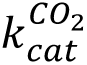 (25.8 s**^−^**^1^; **Figure 1A**) or 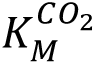 (160–164 µM) (**Supplementary Table 1**), allowing use of immobilized metal affinity chromatography-purifiable C-terminally tagged GWS1B for further work. These GWS1B hexamer (RbcL_6_) carboxylation properties are similar to dimeric *Gallionella* sp. Rubisco (RbcL_2_; 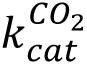 = 27.5 s**^−^**^1^) (2, 39), despite their RbcL sharing only 68% primary sequence identity (**Supplementary Table 1**). Both of these *Gallionella* Rubisco isoforms are approximately 2-fold faster than the model *Rhodospirillum rubrum* Form II Rubisco (**Figure 1A**), and share a common affinity for CO_2_ (149–165 µM) and *S*_C/O_ (9 mol/mol; **Supplementary Table 1**). The exceptionally fast GWS1B does, however, display a heightened O_2_ sensitivity (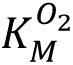 = 98 µM) relative to *R. rubrum* Rubisco (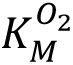 = 159 µM).

**Figure 1.**
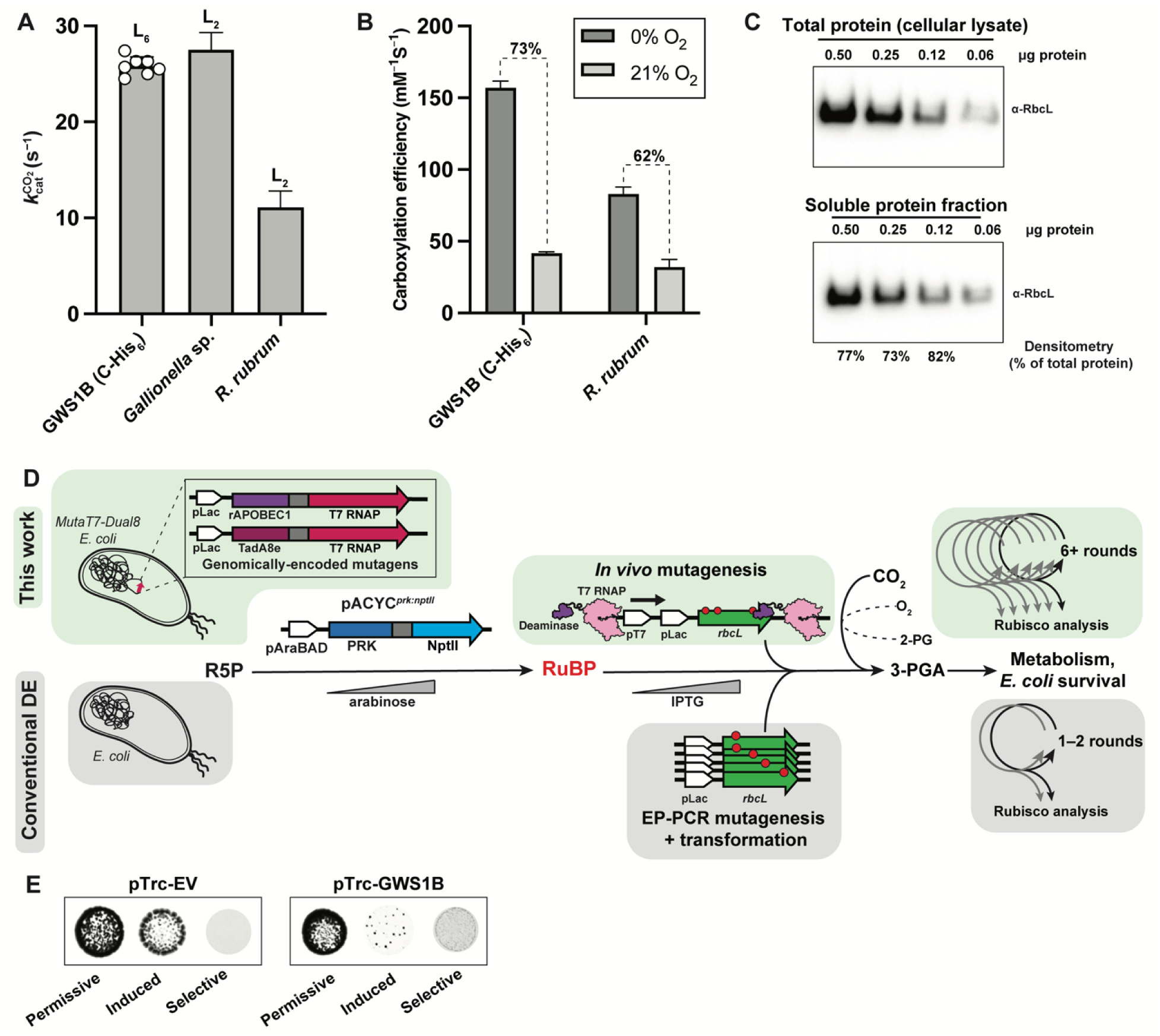
Biochemistry of GWS1B Rubisco and potential for *in vivo* directed evolution in *E. coli*: **A.** Carboxylation rates for C-terminally-tagged GWS1B Rubisco (this work), as well as for *Gallionella* sp. Rubisco (2) and *R. rubrum* Rubisco (48). **B.** Carboxylation efficiencies at 0% O_2_ and 21% O_2_ for GWS1B and *R. rubrum* Rubisco (48). **C.** Western blot analysis of Rubisco levels detected in total protein lysate (top) compared to the soluble protein fraction (bottom) when GWS1B is expressed from the pTrc-GWS1B plasmid, revealing the very high soluble expression of GWS1B. **D.** Traditional Rubisco directed evolution workflow versus targeted mutagenesis-mediated *in vivo* directed evolution workflow. *In vivo* directed evolution is enabled by in-cell, MutaT7-based mutagenesis rather than by extracellular, error-prone PCR (ep-PCR) mutagenesis followed by transformation into *E. coli* cells. MutaT7-based mutagenesis, enabled by a deaminase-T7 RNA polymerase (T7 RNAP) fusion binding to and processing from the T7 RNAP promoter site (pT7) on *rbcL*, generates high levels of downstream mutations in the plasmid DNA and avoids the need for step-wise generation of mutations and transformation, enabling deeper mutagenesis and more rapid progress between rounds of directed evolution. In both workflows, an exogenous *prk* gene is expressed to create a circuit where *E. coli* survival is dependent on Rubisco-catalyzed carboxylation and detoxification of RuBP. R5P = Ribulose-5-phosphate; RuBP = Ribulose-1,5-bisphosphate; 2-PG = 2-phosphoglycolate; 3-PGA = 3-phosphoglycerate. **E.** RDE testing of GWS1B against an empty vector (pTrc-EV) control. Permissive growth media contains no additives, induced growth media contains 0.5 mM IPTG for expression of Rubisco, and selective growth media contains 0.5 mM IPTG, kanamycin at 400 mg/mL, and 0.15% L-arabinose for expression of PRK-NPTII.

This increased sensitivity to O_2_ causes the carboxylation efficiency of GWS1B under anaerobic versus ambient O_2_ conditions to differ by 73%, compared to just a 62% difference for *R. rubrum* Rubisco (**Figure 1B**) (30, 37, 38, 48). These findings emphasize how high Rubisco CO_2_ fixation rates can be achieved via multiple evolutionary routes (*i.e.* are not limited to an isolated region of sequence space) with varying impact on the enzymes’ affinity and specificity for CO_2_ and O_2_.

Alongside GWS1B’s extreme CO_2_ fixation rate and O_2_ sensitivity, GWS1B displays a remarkably high solubility upon expression in *E. coli* (∼77% of the total expressed RbcL; **Figure 1C**). This property could provide a compelling advantage for directed evolution in RDE when aiming to select for kinetic, rather than solubility, enhancing mutants. In prior RDE studies with cyanobacterial Rubisco, where 80–99% of translated RbcL monomers form insoluble aggregates, the majority of higher fitness-conferring RbcL variants identified were those that enhanced RbcL folding and assembly into soluble RbcL_8_RbcS_8_ complexes (21, 23, 25, 27–29, 31–33). Thus, we anticipated that the high solubility of GWS1B would be particularly advantageous, as it would bias selection towards improvements in Rubisco carboxylation kinetics rather than solubility. Moreover, we expected that the high levels of soluble GWS1B expression from the single-copy number bacterial artificial chromosome (BAC) vectors used in MutaT7-driven *in vivo* directed evolution campaigns would be beneficial for successful RDE selection from a single-copy vector (46).

### *In vivo* directed evolution of GWS1B Rubisco using MutaT7-RDE

Conventional Rubisco directed evolution campaigns rely on gene diversification by error-prone (ep)-PCR, followed by library screening using RDE or *rbcL* null photosynthetic hosts (**Figure 1D**, conventional approach) (21–29, 31–33, 48). These campaigns, while occasionally successful in identifying higher carboxylase activity Rubisco catalytic mutants, have suffered from low-mutagenic throughput, owing to the labor-intensive process of ep-PCR-mediated library creation, colony-picking, and analysis. We developed a new Rubisco evolutionary pipeline combining MutaT7 with RDE and next generation sequencing (NGS) to create an *in vivo* Rubisco mutagenesis system suited to repeated rounds of directed evolution in quick progression. Our system utilizes MutaT7-Dual8, a DH10B-derived *E. coli* strain that contains two genomically-encoded mutagenic T7 RNAP fusions that install transition mutations at loci between a T7 RNAP promoter and terminator sequence (42) (**Figure 1D**). MutaT7-Dual8 was transformed with the plasmid pACYC*^prk:nptII^*, enabling arabinose-tunable expression of a phosphoribulokinase–neomycin phosphotransferase fusion (PRK-NPTII) that both phosphorylates the endogenous metabolite ribulose-5-phosphate into ribulose-1,5-bisphosphate (RuBP, the sugar substrate of Rubisco) and confers kanamycin resistance. As RuBP is a foreign and toxic metabolite to *E. coli*, cell survival can be made dependent on Rubisco-mediated RuBP clearance (23, 33, 48). Such RDE screens include selection on kanamycin to avoid the growth of any false positives arising from mutations in pACYC*^prk:nptII^* that might impair PRK, and hence toxic RuBP, production (**Figure 1D**).

As observed previously, the production of Rubisco in *E. coli* can itself impair cell viability (30, 49). Indeed, high level expression of GWS1B in DH10B impairs colony growth (**Figure 1E**, ‘induced’). Still, in pACYC*^prk:nptII^* transformed DH10B (here termed DH10B-RDE selection) we observed that cell growth was dependent on GWS1B expression under 0.15% w/v arabinose PRK-NPTII induction (**Figure 1E**, ‘selective’). These findings confirmed the suitability of GWS1B for RDE-mediated selection of mutants that enhance cell fitness during PRK-NPTII induction.

We thus transformed MutaT7-Dual8 containing pACYC*^prk:nptII^*with a MutaT7-targeting BAC vector encoding polyhistidine-tagged GWS1B (BAC-GWS1B). We named this mutagenesis-selection system, containing all the necessary components for *in vivo* directed evolution of Rubisco, MutaT7-RDE (**Figure 1D**, this work).

Within just 30 days, BAC-GWS1B was subjected to six rounds of evolution in MutaT7-RDE under incremental increases in arabinose-induced PRK-NPTII selection pressure (**Figure 2A**). Each round consisted of a 24-hour MutaT7-induced continual mutagenesis period in liquid culture, before plating on selective media and incubating in air supplemented with 10% v/v CO_2_ for three days. The resulting *E. coli* colonies were harvested and the BAC-GWS1B and pACYC*^prk:nptII^* plasmids were isolated for short read next-generation sequencing (NGS). The co-purified plasmids were also *Sfi*I-digested (cutting only pACYC*^prk:nptII^*) and the BAC-GWS1B plasmids retransformed into either fresh MutaT7-RDE cells for the next round of MutaT7-RDE selection, or into DH10B-RDE for re-screening and whole-plasmid sequencing of any faster-growing ‘winner’ clonal isolates (**Figure 2A**).

**Figure 2.**
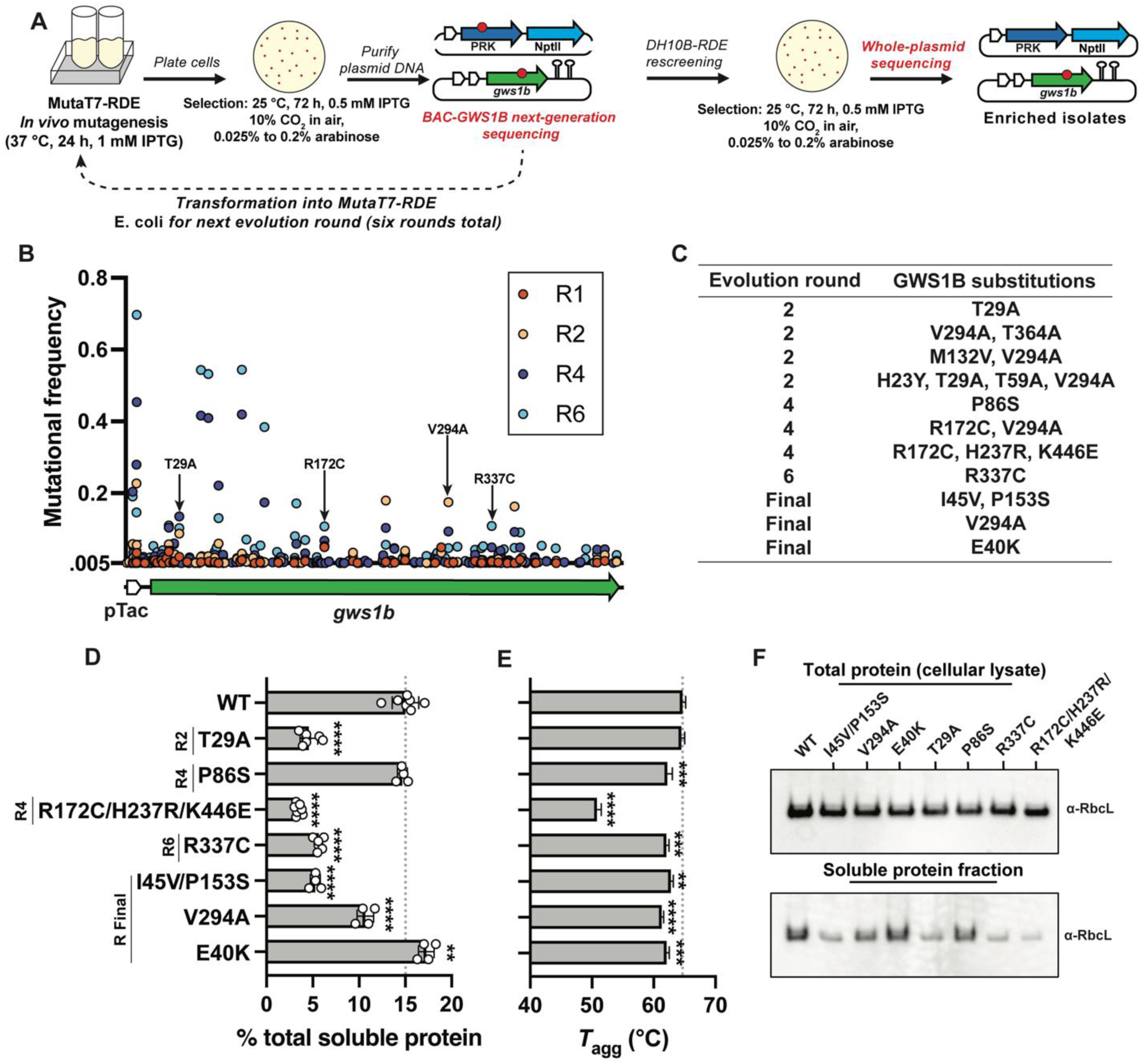
Identification and biochemical characterization of GWS1B variants: **A.** Schematic of MutaT7-RDE workflow. **B.** Map of the promoter and GWS1B-coding region of the BAC evolution plasmid showing mutational frequency at the DNA base level during sequenced rounds of evolution. Non-synonymous mutations from clonal isolates are indicated with arrows **C.** Non-synonymous mutations found in clonal isolates of GWS1B in rounds 1, 2, 4, 6, and final rescreening round of evolution. **D.** Soluble, folded Rubisco content of selected variants in *E. coli* quantified by [^14^C]-CABP binding (50). Data are the means and standard deviations of three replicates with significance shown relative to wild-type GWS1B as analyzed by one-way ANOVA. **, *p* ≤.005; ***, *p* ≤.0005; ****, *p* ≤.0001. **E.** Aggregation temperature (*T*_agg_) of selected variants. Significance testing and labels are consistent with (**D**). **F.** SDS-PAGE immunoblots of GWS1B RbcL levels in the soluble and total protein fractions upon expression of GWS1B variants in *E. coli*.

The short-read NGS data revealed that the high-frequency (above detection threshold) mutations that accrued in BAC-GWS1B during MutaT7-RDE selection were highly enriched between the T7 promoter and terminator regions (**Figure 2B** and **Supplementary Figure 1**) (42), including in the tac promoter sequence driving GWS1B expression (**Supplementary Table 2**). As expected, transition mutations accounted for all non-synonymous mutations in the *gws1b* gene. Encouragingly, mutational frequencies at over 20 specific sites in *gws1b* showed substantial variation between evolutionary rounds two and four and generally continued to enrich with increasing selection pressure into the sixth and final DH10B-RDE selection round (**Supplementary Table 2**). Comparative sequence alignment of GWS1B with other members of the Rubisco family (**Supplementary Figure 2**) showed that the selected GWS1B mutations occurred in poorly conserved regions that plausibly are more tolerant to mutational change, and hence are more readily accessible during MutaT7-RDE selection.

Whole-plasmid sequencing from DH10B-RDE rescreened clonal isolates following evolution rounds 2, 4, and 6 identified multiple non-synonymous mutations in *gws1b* (**Figure 2C**). In addition, many synonymous mutations in *gws1b* and mutations in the promoter and plasmid replication elements were detected (**Supplementary Table 3**). Whether these extraneous mutations influenced DH10B-RDE fitness through artefactual impacts on GWS1B expression or BAC-GWS1B copy number remain untested. We focused our attention instead on 11 *gws1b* mutant genes identified via isolate screening (**Figure 2C**), with each cloned into a clean pTrc backbone to avoid any vector mutation artifacts, expressed in DH10B *E. coli*, and then purified for biochemical analysis.

### Evaluating the speed, solubility, and stability of GWS1B Rubisco variants

Initial comparisons of relative carboxylase activities measured via spectrophotometry (2, 50) indicated that five of the purified mutant GWS1B enzymes shared wild-type like carboxylation rates, three had 2–10-fold lower rates and three displayed little or no measurable activity (**Supplementary Figure 3**). How the three apparently inactive variants supported growth during DH10B-RDE rescreening remains unclear, and could even relate to a loss of activity during the purification process. We do note, however, that directed evolution campaigns, including RDE-based selection campaigns, often encounter false positives (24, 26, 33, 49) – for example, owing to plasmid instability (33) or transposon insertion in *prk* (22, 23, 49). Fusion of PRK to NPTII should eliminate false positives owing to transposon insertion or plasmid instability, but does not protect against ablation of PRK function via a point mutation. Indeed, several point mutations in pACYC*^prk:nptII^*were identified by NGS in each round of MutaT7-RDE selection (**Supplementary Figure 4**). Some of these mutations may reduce or destroy the ribulose-5-phosphate phosphorylation activity of PRK-NPTII while maintaining kanamycin resistance. The acquisition of comparable mutations in pACYC*^prk:nptII^*during DH10B-RDE rescreening may have allowed growth of some escape mutants even with reduced GWS1B Rubisco function.

We proceeded with the six GWS1B variants that displayed wild-type or better carboxylation activity (**Supplementary Figure 3**). We first sought to assess whether these variants were selected due to improved GWS1B solubility, rather than improved kinetic features. We quantified the percentage of soluble, folded GWS1B relative to total cellular protein using a ^14^C-2-Carboxyarabinitol-1,5-bisphosphate (^14^C-CABP) binding assay (50). This assay revealed that the P86S and E40K GWS1B variants, selected in the fourth and final selection rounds, respectively, shared wild-type levels of GWS1B production in DH10B *E. coli*. All three enzymes accumulated to ∼15% w/w of the soluble cellular protein (**Figure 2D**). By contrast, the other variants displayed 20–80% reductions in soluble, folded GWS1B production, despite the temperature of aggregation (*T*_agg_) for all of these variants appearing to retain near wild-type levels of stability (**Figure 2E**). Moreover, all six successfully formed the RbcL_6_ holoenzyme as assessed using native PAGE (**Supplementary Figure 5**). We also assessed the lower-activity triple mutant GWS1B R172C/H237R/K446E using these methods. There was noticeable precipitation of this variant during purification – possibly explaining the inability of the initial spectrophotometric assays to detect activity (**Supplementary Figure 3**). We indeed observed that the triple mutant led to poor production of soluble GWS1B and lowered the *T*_agg_ > 10 °C *in vitro*, likely limiting the amount of RbcL_6_ complexes detectable by ^14^C-CABP binding (**Figures 2D** and **2E**).

Additional SDS-PAGE and immunoblotting analyses showed that, while wild-type equivalent levels of each GWS1B RbcL variant were translated in DH10B *E. coli* (lysate sample; **Figure 2F**), the proportion of produced GWS1B that was soluble in the cytosol differed between the variants (soluble sample; **Figure 2F**). Notably, the relative levels of soluble RbcL detected by blotting correlated well with the proportion of functional enzyme quantified by ^14^C-CABP binding (**Figure 2D**), suggesting that the soluble RbcL detected accurately reflects the fraction stably assembled as a functional RbcL_6_ holoenzyme. In sum, while none of the selected GWS1B variants had any discernable impact on GWS1B RbcL translation, the majority of substitutions quite substantially limited the folding and assembly of RbcL_6_ holoenzyme.

Together, these observations indicated the GWS1B variants selected in our platform did not arise from increases in solubility, thus motivating their kinetic evaluation..

### Comprehensive kinetic evaluation of evolved GWS1B Rubisco variants

We performed a full kinetic analysis of the GWS1B variants T29A, P86S, R337C, I45V/P153S, V294A, and E40K, as well as the triple mutant R172C/H237R/K446E using radiometric carboxylation activity assays (**Figures 3A–G** and **Supplementary Table 4**). The carboxylation rate of all variants remained similar to or decreased relative to wild-type GWS1B (**Figure 3A**), consistent with the spectrophotometric estimate of carboxylation rate (**Supplementary Figure 3**). This result was expected, owing to the already extremely high carboxylation rate of wild-type GWS1B that limits selection pressure on this parameter.

**Figure 3.**
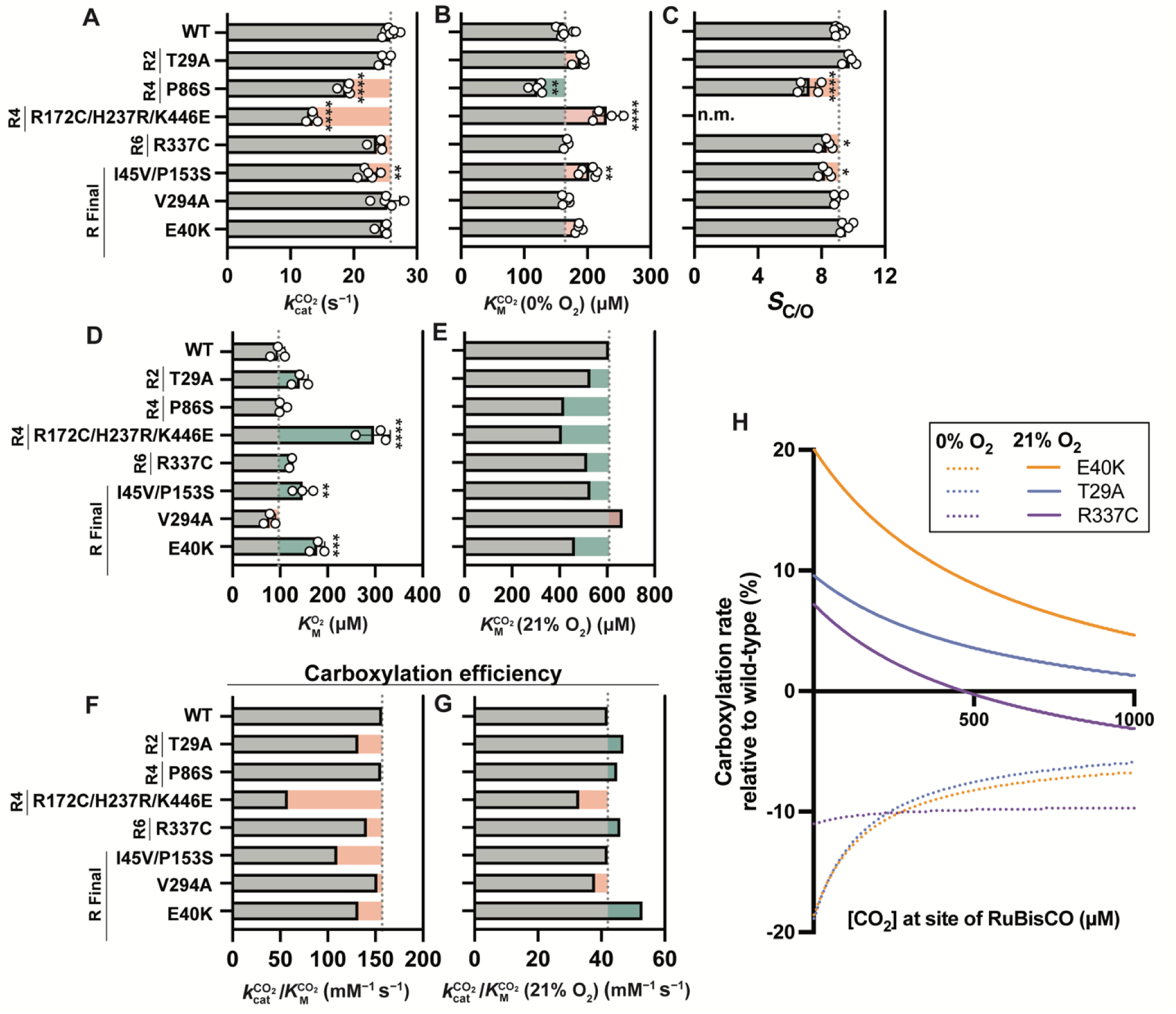
Kinetic characterization of GWS1B variants: **A.** Carboxylation rate. **B.** Affinity for CO_2._ **C.** Specificity factor. **D.** Affinity for O_2_. Asterisks indicate *p*-value as calculated by one-way ANOVA followed by a post-hoc Tukey test. *, *p* ≤.05; **, *p* ≤.005; ***, *p* ≤.0005; ****, *p* ≤.0001. **E.** Affinity for CO_2_in air. **F.** Carboxylation efficiency at 0% O_2_. **G.** Carboxylation efficiency at 21% O_2_. **H.** Carboxylation rates of GWS1B E40K, GWS1B T29A, and GWS1B R337C relative to wild-type GWS1B at 0% or 21% O_2_ as a function of CO_2_ concentration.

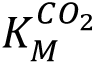 measured at 0% O_2_ increased for all but one variant (P86S; **Figure 3B**). *S*_C/O_ remained comparable or slightly decreased relative to wild-type GWS1B (**Figure 3C**).

Critically, however, 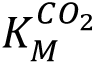 at 0% O_2_ is an irrelevant parameter in the context of aerobic RDE selection. That is, we evolved the normally near-anaerobic GWS1B enzyme under the unnatural selection pressure of ambient O_2_ concentrations with the hypothesis that GWS1B variants with improved fitness would show gains in resistance to O_2_ inhibition. To test this hypothesis, we derived the apparent *K_M_* for O_2_ 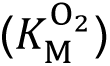 for each GWS1B variant (**Figure 3D** and **Supplementary Figure 6**). In almost all instances, the selected variants reduced the sensitivity of GWS1B to O_2_ inhibition (i.e., the variants increased 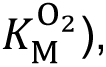 with only V294A increasing the enzyme’s apparent affinity for O_2_ (**Figure 3D**). Upon accounting for these changes in O_2_ affinity, the 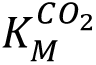 under ambient oxygen [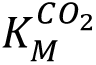(21% O_2_)] for each enzyme showed that all the selected variants, except V294A, had higher than wild-type aerobic affinities for CO_2_ (**Figure 3E** and Supplementary Table 4).

Taken together, these data reveal that while the anaerobic carboxylation efficiencies of each evolved GWS1B variant do not exceed wild-type (**Figure 3F**), under atmospheric O_2_ the carboxylation efficiency of the T29A, R337C, and E40K GWS1B variants are improved 11%, 8%, and 25%, respectively (**Figure 3G**). The consequence is that, relative to wild-type GWS1B, these three selected variants can support higher rates of CO_2_ fixation in ambient O_2_ environments (**Figure 3H**; solid lines) but not in an anaerobic setting (**Figure 3H**; dashed lines).

Particularly noteworthy is the broad CO_2_ range under which the identified substitutions would benefit GWS1B aerobic catalysis, with even the weaker performing R337C substitution supporting faster than wild-type carboxylation rates up to ∼500 µM CO_2_ (**Figure 3H**), a concentration equivalent to that naturally achievable in cyanobacterial carboxysomes during CO_2_-concentrating mechanism (CCM) induction (51).

It is unclear how the V294A, I45V/P153S and R172C/H237R/K446E variants were selected given their impaired kinetic fitness. While in terms of altered affinities for CO_2_ and O_2_, the triple mutant showed the greatest improvement [247% higher 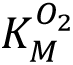; 33% lower 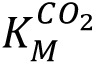(21%O_2_)], its 2-fold lower 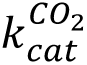 impaired aerobic carboxylation efficiency by 23% relative to wild-type GWS1B (**Supplementary Table 4**). Considering the cellular toxicity associated with GWS1B production in *E. coli* (**Figure 1E**), one possibility is that the lower solubility of these three variants (**Figures 2D** and **2F**), may somehow have provided a fitness advantage that improved MutaT7-RDE and DH10B-RDE growth during selection. Indeed, *E. coli* expressing GWS1B V294A or I45V/P153S grew more quickly than *E. coli* expressing wild-type GWS1B (**Supplementary Figure 7**).

### Selected substitutions in GWS1B cluster near the active site and at protein**–** protein interfaces

Structural analysis of the carboxylase-enhanced GWS1B variants show the N-domain substitution T29A is located 20 Å away from the active site behind an α-helix, with N-domain E40K 15.4 Å away from the active site (**Figure 4A**). The C-domain R337C substitution is located very distant from any active site on the surface distal to both dimer and hexamer interfaces (**Figure 4B**).

**Figure 4.**
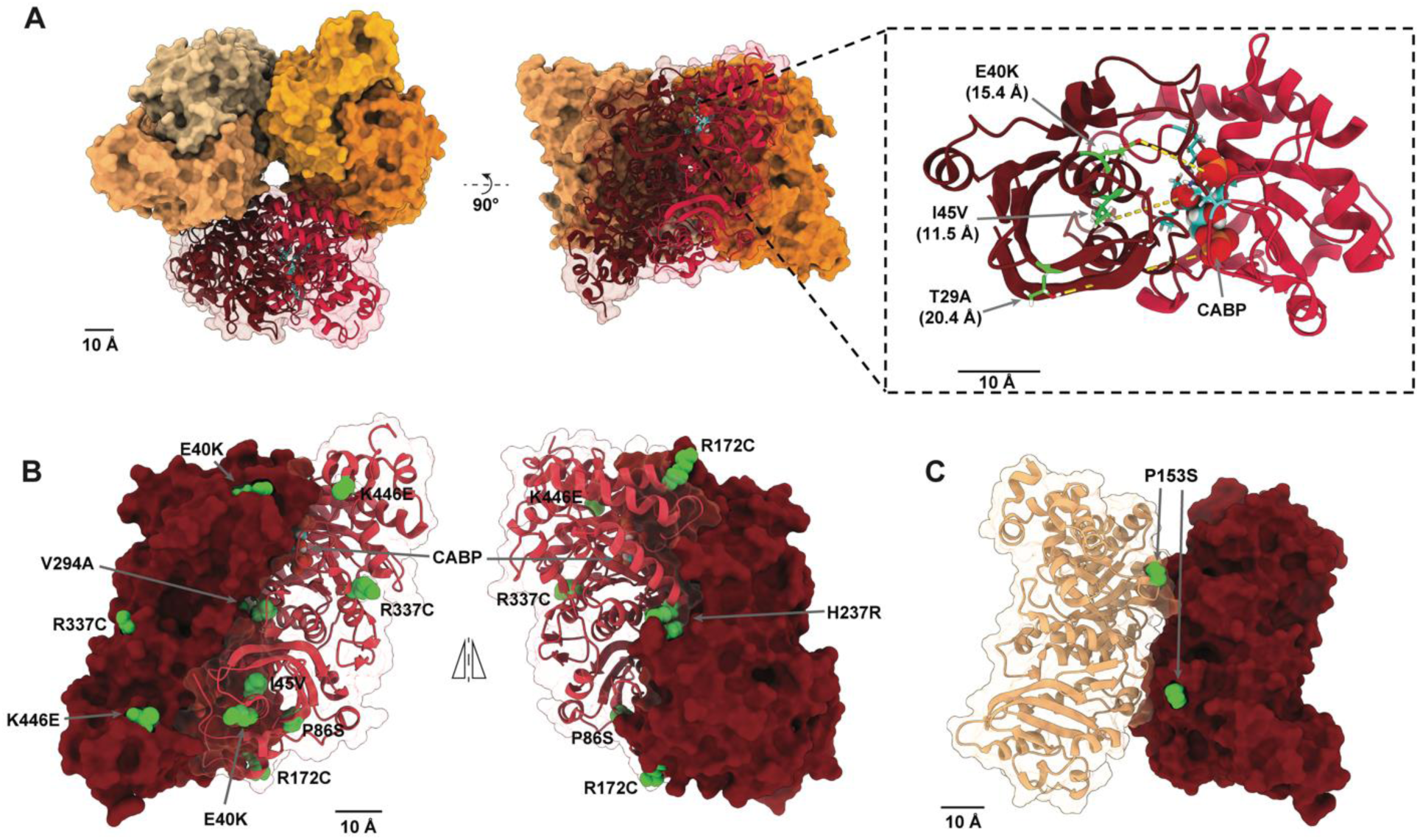
Mapping evolved variants onto GWS1B: **A.** Substitutions T29A, E40K, and I45V are positioned near the GWS1B active site. PDBID: 5C2G (37). **B.** E40K, I45V, P86S, R172C, H237R, V294A, and K446E occur at the dimer interface. R337C is surface-exposed. **C.** P153S occurs at the hexamer interface of GWS1B.

Similarly to E40K, whose position near the active site also positions it near the RbcL– RbcL interface, the solubility-influencing substitutions P86S, R172C, H237R, V294A, and K446E all locate to the RbcL–RbcL interface (**Figure 4B**), and P153S lies at the interface between adjoining RbcL_2_ dimers (**Figure 4C**). These observations suggest the observed solubility impediments may not arise from changes in chaperonin-assisted folding of the GWS1B RbcL monomers, but rather that the amino acid substitutions at the surface of the RbcL–RbcL interface impede RbcL_2_ and RbcL_6_ oligomeric assembly. The kinetics of these variants show that the substitutions inflict long range structural rearrangements on the active site architecture that, in most instances, unwantedly impaired catalytic performance (**Supplementary Table 4**).

Mapping the location of the selected GWS1B RbcL substitutions revealed that they typically co-located to the respective functional region (active site or interface) of the model Form II *R. rubrum* Rubisco and model Form I plant *N. tabacum* Rubisco (**Supplementary Table 5**). This observation suggests substitutions at these corresponding RbcL residues may also impart favorable kinetic properties across the Rubisco family. Indeed, when compared to the *R. rubrum* Rubisco deep mutational scanning analysis of Prywes et al. (2025) (30), 50% of the selected GWS1B mutations occurred at loci where fitness was improved in *R. rubrum* Rubisco (**Supplementary Table 5**). This outcome is striking when one considers that only 0.14% of substitutions in the *R. rubrum* Rubisco survey were differentiated as fitness-improving. Taken together, the improvements we observed in Rubisco catalysis in GWS1B reveal multiple pathways to O_2_ resistance, and support the notion that laboratory-based *in vivo* directed evolution can push Rubisco beyond existing natural catalytic boundaries.

## Discussion

In this work, we introduce an *in vivo* directed evolution platform for Rubisco enzymes and use it to evolve Rubisco from *Gallionellaceae*, yielding variants with improved carboxylation efficiency and rates in air. This system builds upon years of foundational work on Rubisco evolution by accelerating the duration of evolution rounds, reducing researcher intervention, and deeply sampling Rubisco sequence space. The success of this evolution campaign is both a proof-of-concept for *in vivo* Rubisco directed evolution and a demonstration of exceeding the known limits of carboxylation capacity for a Form II Rubisco, a feat so far untenable owing to the inherently fast speed and high *E. coli* solubility exhibited by the primitive homo-oligomeric enzymes in this lineage (30, 48).

Exerting an oxygenic selection pressure on a Rubisco isoform isolated from a low-oxygen environment yielded improvements to 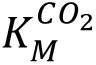(21% O_2_) and 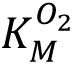, leading to a substantial net increase in carboxylation efficiency in air for this normally highly O_2_-sensitive Rubisco. The GWS1B variants identified here demonstrate that anaerobic and ambient carboxylation efficiencies are not intrinsically linked. Thus, going forward, the effects of O_2_ on Rubisco activity must be considered when undertaking any Rubisco engineering campaign to fully grasp and analyze changes in catalytic fitness, particularly for C3 plant Rubisco that contend for available CO_2_ in an environment rich in dissolved O_2_.

The mechanism by which the selected mutations provide O_2_ resistance is not yet clear. The substitutions T29A and E40K, which enhance ambient carboxylation efficiency, are within 20 Å of the active site, possibly creating subtle structural changes that benefit the affinity for CO_2_ over O_2_. Seven selected substitutions that lie at either the dimer or hexamer interface of

GWS1B altered 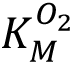, though not always resulting in an increase in carboxylation efficiency in an ambient atmosphere. This observation is not the first instance of amino acid substitutions at Rubisco protein–protein interfaces affecting relative affinities for CO_2_ and O_2_. Selected substitutions clustered at the interface between the large and small Rubisco subunits during a previous directed evolution of the Form I *Thermosynechococcus elongatus* Rubisco, causing an increase in carboxylation efficiency in air (23). Designed structural substitutions that transition GWS1B from hexamer to dimer also impacted catalysis, increasing both *S*_C/O_ and, to a larger extent, 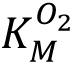, albeit at the cost of decreasing 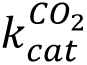 and carboxylation efficiency (38). A recent deep mutational scan of Form II *R. rubrum* Rubisco identified only a handful of single amino acid substitutions, all at the dimer interface, that enhanced affinity for CO_2_ under anoxia, albeit accompanied by substantial decreases in 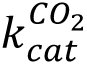 or carboxylation efficiency, and with undetermined impact on 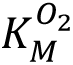 (30). Hence, there is accumulating evidence, including our work here, that residues distant from the active site at RbcL among differing Rubisco lineages have a pervasive, and sometimes beneficial, impact on catalysis. Moreover, for GWS1B, the T29A, E40K, and R337C substitutions provide additional examples that subvert the tradeoff between CO_2_ affinity and carboxylation rate that has debilitated the aforementioned variants in other Form II Rubisco. They also emphasize the importance of scrutinizing how mutations impact the O_2_ sensitivity of a Rubisco. In the absence of such studies, essential insight into how the enzyme’s carboxylation rate under ambient O_2_ is impacted by novel substitutions is lacking. As indicated above, such understanding is critical when attempting to simulate performance in an aerobic photosynthetic environment, such as a leaf.

In conclusion, we deployed a novel *in vivo* directed evolution system to expand the catalytic limits of Rubisco beyond what has been observed in nature, focusing on minimizing inhibitory oxygenation while stimulating carboxylation. Future use of MutaT7-RDE, as well as orthogonal protein engineering approaches, will continue to elucidate the Rubisco sequence-structure-function landscape. Efficient Rubisco enzymes engineered in the laboratory will provide key insights into the fundamental mechanisms of carboxylation and oxygenation and can be utilized as essential starting points to improve photosynthesis.

## Materials and Methods

### MutaT7-RDE workflow

To generate the MutaT7-RDE strain, a single colony of BAC-GWS1B and pACYC*^prk:nptII^*^-^ transformed Dual8 *E. coli* (42) were grown on agar-solidified Luria broth (LB) containing 100 µg/mL ampicillin and 35 µg/mL chloramphenicol (LB-AC). A single colony was inoculated into 5 mL of LB-AC containing 1 mM isopropyl β-D-1-thiogalactopyranoside (IPTG) to induce MutaT7 mutagenesis. After incubating at 37 °C and 180 rpm for 24 h, a 0.1 µL aliquot of this culture was plated on LB-AC agar containing 50 µg/mL triphenyl tetrazolium chloride (TTC; Sigma-Aldrich) to visualize colony forming units (cfu) per volume of the culture. 250 µL aliquots of the remaining culture (∼5–7 × 10^5^ cfu) were plated on selective LB (LB-AC agar plates with 400 µg/mL kanamycin, 0.5 mM IPTG, and 50 µg/mL TTC) containing between 0.025–0.2% w/v L-arabinose (Sigma-Aldrich) to modulate PRK-NPTII production. L-Arabinose concentrations were incrementally increased by 0.025% w/v between rounds 1 to 4, and then increased to 0.15% w/v (round 5) and 0.2% w/v (round 6). The plates were incubated at ∼25 °C in a commercial pre-mixed high CO_2_ atmosphere (10:21:69% v/v CO_2_:O_2_:N_2_) until colony growth was observed (∼72 h). The resulting colonies were harvested by scraping into 5–10 mL LB, centrifuged for 10 min at 5,000 × *g*, and the BAC-GWS1B and pACYC*^prk:nptII^*plasmids were co-purified (Wizard® *Plus* SV Miniprep kit; Promega) for NGS or further MutaT7-RDE re-screening.

For NGS, the plasmids were diluted to 20 ng/µL for Nextera DNA Flex kit (Illumina) preparation before sequencing in duplicate using a 150 paired-end kit on an Element Biosciences AVITI sequencer. For MutaT7-RDE rescreening, 1 µg of the co-purified plasmids were digested with *Sfi*I (New England Biolabs) for 2 h at 50 °C, re-purified (Wizard® SV PCR clean up kit; Promega) and eluted in 15 µL H_2_O. After electroporating 5 µL into MutaT7-RDE, the cells were recovered in 1 mL LB for 1 h at 37 °C/180 rpm before plating 1 µL on LB-AC to assess transformation efficiency and diluting the remaining cells into 4 mL LB-AC with 1 mM IPTG to initiate the next round of MutaT7 mutagenesis.

For non-mutagenic rescreening, the purified *Sfi*I-digested plasmid was electroporated into pACYC*^prk:nptII^*-transformed DH10B (DH10B-RDE) *E. coli* and plated on selective LB containing 0.2% w/v L-arabinose and incubated at ∼25 °C under 10% v/v CO_2_ for 3–5 d. The fastest-growing colonies were individually grown in LB-AC, and then the plasmids purified and whole-plasmid sequenced (Plasmidsaurus).

To verify variants from the later rounds of evolution, *rbcL* libraries from rounds 4 and 6 were PCR-amplified using the primers GWS1Bfor (5′-ATATCCATGGACCAAAGTAATCGTTACGCCGAC-3′) and GWS1Brev (5′-AATATGAGCTCCTATTCGCCATTCAGGCTGCGTTAATTAAACCAC-3′) and cloned as a *Nco*I-*Sac*I fragment into a pTrc vector. The fastest-growing colonies upon re-screening were picked and analyzed as variants from the “final” round of evolution.

### Next-generation sequencing analysis

Output fastQ files were aligned to BAC-GWS1B and pACYC*^prk:nptII^*using the Burrows-Wheeler Aligner (BWA) (52). Alignments were down-sampled to 10% to reduce file size, then were indexed, sorted, and a pileup file was generated using Samtools (53). Non-reference base frequencies at each position in the reference plasmid were calculated using Samtools and a custom Perl script. Mutational scatter plots were generated from these tables using Prism.

Further analysis of high-percentage SNPs was completed with Geneious Prime. Mutations at 0.10% or higher frequency (determined using a read quality score of 30) in BAC-GWS1B that were present in duplicate sequencing of at least one round of evolution are listed in **Supplementary Table 2**. Full scripts for NGS processing are available at https://github.com/julielmcdonald/GWS1B.

### Rubisco purification

Mutant and wild-type GWS1B *rbcL* genes were PCR-amplified using the primers GWS1Bfor and GWS1Brev, cloned as a *Nco*I-*Sac*I fragment into a pTrc vector, and transformed into DH10B *E. coli*. Large (1 L) or small (10–20 mL) LB containing 100 µg/mL ampicillin (LB-A) cultures were grown for Rubisco purification and high-resolution kinetic analyses, respectively. Cultures were grown at 37°C at 180 rpm until an OD_600_ of 0.6–0.8 was attained, before inducing GWS1B expression with 0.5 mM IPTG at 23 °C for 16 h and then centrifuging for 10 min at 8000 × *g* and 4 °C. Cell pellets were then snap-frozen in liquid N_2_ for storage at −80 °C.

For Rubisco purification, the stored cell pellets were resuspended in 20 mL ice cold buffer [20 mM HEPES, pH 8.0, 50 mM NaCl, 1 mM phenylmethylsulfonyl fluoride (PMSF; Amersco), 5 mM DL-Dithiothreitol (DTT; Sigma-Aldrich) and 1 µL of DNase (Sigma-Aldrich)] then lysed using a needle-tipped sonicator (50% amplitude with three 20 s on/off cycles) on ice.

Following centrifugation (20,000 × *g* for 20 min at 4°C), the soluble protein was filtered through a 0.22 µm filter before immobilized metal affinity chromatography purification using a 1 mL HisPur™ Cobalt Resin (Thermo Fisher Scientific) column equilibrated with column buffer (20 mM HEPES and 50 mM NaCl at pH 8.0). After washing with 5 mL column buffer with 20 mM imidazole (Alfa Aesar), GWS1B proteins were eluted with 2 mL column buffer containing 500 mM imidazole. The eluted protein was dialyzed at 4 °C o/n against column buffer, concentrated using an Amicon^®^ Ultra Centrifugal Filter (Sigma-Aldrich) with a 100 kDa molecular weight cutoff, and then snap-frozen for future use. Purity and oligomeric state were evaluated using Native PAGE (**Supplementary Figure 5**), and GWS1B concentration was calculated from NanoDrop 280 nm absorbance using an extinction coefficient of 63175 M^−1^ cm^−1^ for GWS1B. Purified Rubisco were used to quantify *S*_C/O_ using the [1-^3^H]-RuBP consumption assay (54).

### Spectrophotometric measurement of Rubisco activity

Carboxylation rates for each purified GWS1B enzyme were measured using the enzyme-coupled nicotinamide adenine dinucleotide (NAD^+^/NADH) reduction spectrophotometric assay in 96-well microtiter plates, as previously described (2, 50). Purified GWS1B proteins (1 mg/mL) were activated in 100 mM EPPS-NaOH at pH 7.6 containing 10 mM MgCl_2_ and 10 mM NaHCO_3_ before diluting 1:500 into 0.2 mL reactions containing 10 mM MgCl_2_ and 10 mM NaHCO_3_ and initiating with 5 µL 20 mM RuBP (50). Assays were performed in triplicate at 30 °C, tracing the decrease in absorbance at 340 nm over 20 min at 53 s intervals. The slope of the linear decline at 340 nm over time was used to calculate the Rubisco carboxylation rate (s^-1^) (50).

### GWS1B Rubisco content and kinetic characterization

Bacterial pellets were sonicated on ice (30 s at 60% amplitude) in 0.5–1.0 mL assay buffer (50 mM EPPS-NaOH at pH 7.8 containing 15 mM MgCl_2_, 1 mM EDTA, 1 mM PMSF, and 5 mM DTT), then centrifuged (20,000 × *g* for 2 min at 2 °C) and aliquots of the soluble protein were used for quantifying protein content (Pierce™ Bradford Protein Assay Kit against bovine serum albumin), Rubisco content measurements, ^14^CO_2_ fixation activity assays, and SDS-PAGE analyses. Samples were pre-activated with 25 mM NaHCO_3_ for 10–20 min at 25 °C before quantifying Rubisco content by ^14^C-CABP binding, as previously described (50, 55). Measures of Rubisco concentration as a % (w/w) of total cell soluble protein assumed a molecular mass of 50 kDa for GWS1B RbcL.^14^CO_2_-fixation assays to measure 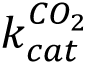, 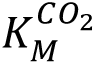 and 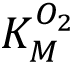 at 25°C were undertaken as previously described (23), with the following additional considerations for GW1SB: Following activation of Rubisco with 20 mM NaH^14^CO_3_, the samples were diluted 50-fold into assays containing six incremental ^14^CO_2_ concentrations between 0–360 µM, 0–440 µM, or 0–540 µM in reactions preequilibrated with atmospheres containing 0% (anaerobic), 10%, or 20% v/v O_2_ in N_2_, respectively (**Supplementary Figure 6**).

### SDS-PAGE, Western Blotting, and Native PAGE

For SDS-PAGE and immunoblotting, samples of total cellular protein following sonication (lysate), as well as the soluble protein following centrifugation (soluble), were processed for SDS-PAGE separation through NuPAGE™ Bis-Tris Mini Protein Gels, 4–12% (Thermo Fisher Scientific), as previously described (23). Sample loading was normalized to soluble protein content (with matching volume of lysate loaded per sample), and the separated proteins either visualized by Coomassie staining or transferred onto nitrocellulose membranes (Hybond C, APBiotech) using an Xcell transfer cell (Novex), according to the manufacturer’s specifications. Immunoblotting was performed as previously described (56), and GWS1B RbcL contents were compared by probing with an antibody to *R. rubrum* Rubisco (56). Immunoreactive GWS1B RbcL was visualized using the Clarity Western ECL substrate (Bio-Rad). The fluorescence signal was imaged and signal densitometry quantified using a ChemiDoc MP Imaging System (Bio-Rad).

For native PAGE, purified GWS1B proteins were thawed on ice and diluted to 1 µg/mL in 20 mM HEPES and 50 mM NaCl at pH 8.0. 1 µg of protein was loaded onto a Novex™ Tris-Glycine 4–12% Mini Protein Gel (Thermo Fisher Scientific). The gel was run in NativePAGE™ Running Buffer (Thermo Fisher Scientific) at 120 V for 2.5 h at 4 °C, then stained with Coomassie InstantBlue® Protein Stain (Abcam). 1 µg of bovine thyroglobulin (Sigma-Aldrich) and 1 µg of bovine serum albumin (Sigma-Aldrich) were used as molecular weight markers.

### Growth curves

pTrc-GWS1B plasmids were transformed into DH10B chemically competent *E. coli* (New England Biolabs). A single colony was used to inoculate 5 mL of LB-A and was incubated overnight at 37 °C and 180 rpm. This culture was used to inoculate 200 µL of LB-A to an OD of 0.005 in a 96-well plate. OD was measured at 600 nm for 16 h +/-IPTG in an Agilent BioTek Synergy HTX plate reader. Data reported are means of normalized triplicate growth curves.

### DH10B-RDE spot testing

Colonies of pTrc-GWS1B and pACYC*^prk:nptII^* co-transformed DH10B *E. coli* (DH10B-RDE) were grown on LB-AC containing 50 µg/mL TTC. A colony was used to inoculate 1 mL of LB-AC, which was then incubated at 37 °C and 180 rpm until an OD_600_ of ∼0.2. The resulting culture was diluted with LB to an equivalent OD_600_ of 1 x 10^-4^ and 20 µL was spotted onto an LB-AC plate containing 50 µg/mL TTC, 0.5 mM IPTG, 400 µg/mL kanamycin and incremental concentrations (0–0.3% w/v) of L-arabinose. The plates were incubated at ∼25 °C in air supplemented with 2% v/v CO_2_ and colony growth was monitored over 10 d.

### Data, Materials, and Software Availability

All study data are included in the article and/or SI Appendix. Plasmids pACYC^prk:nptII^, BAC-GWS1B and pTrc-GWS1B are available on Addgene. MutaT7-Dual8 is available from the Belgian Coordinated Collections of Microorganisms, catalog LMBP 13434.

## Acknowledgements

This work was supported by the National Science Foundation’s Division of Molecular and Cellular Biosciences (EAGER Grant 2244770) and the National Institutes of Health/National Institute of General Medical Sciences Grant 1R35GM136354 to M.D.S., the Abdul Latif Jameel Water and Food Systems Lab Grand Challenge Grant to M.D.S., R.H.W., and S.W., a generous gift to MIT from an anonymous donor, the Martin Family Society Fellowship for Sustainability to J.L.M, and the MIT Robert J Sibley, Future of Science, and BroadIgnite Fellowships to A.A.M. We extend our gratitude to the MIT BioMicroCenter staff for their assistance with high-throughput sequencing experiments. We thank Dr. Jiacheng Lin for his helpful comments and editing of the manuscript.

## Notes

### Competing Interest Statement

The authors have declared no competing interest.

### Summary of Updates

RuBisCO -> Rubisco. Minor edits to text in Discussion. Values in Fig. 1A and p. 6 from Davidi et al. (2020) were updated to reflect a published correction (Tsai et al. (2025)). One co-author was added.

