## Supplementary Data for "*In Vivo* Directed Evolution of an Ultra-Fast Rubisco from a Semi-Anaerobic Environment Imparts Oxygen Resistance"

**This PDF file includes:**

Tables S3, S6
Figures S1 to S7
SI References

**Other supporting materials for this manuscript include the following:**

Tables S1, S2, S4, S5

**Supplementary Table 1.** Comprehensive kinetic analysis for wild-type GWS1B (C- and N-terminally-tagged, untagged), *Gallionella* sp. Rubisco, and *Rhodospirillum rubrum* Rubisco (see Excel file).

**Supplementary Table 2.** Mutants from next-generation sequencing (see Excel file).

**Supplementary Table 3.** Synonymous and other substitutions from isolate sequencing.

| Evolution round | GWS1B substitutions | Tac promoter mutations | Silent GWS1B variants | Other substitutions |
|---|---|---|---|---|
| 2 | T29A | - | I18 | repE E101K, K107I |
| 2 | V294A, T364A | t477c | I232, G359 | repE K107I, H120R, Q196R, |
| 2 | M132V, V294A | a478g | E40, I232, G359 | repE K107I, H120R, Q196R; sopA P345Q; sopB A229T |
| 2 | H23Y, T29A, T59A, V294A | a478g | I232, G359 | repE K107I, H120R, Q196R; sopA P345Q |
| 4 | P86S | c468t | T59, F73 | repE E106K, K107I |
| 4 | R172C, V294A | t477c | I232, G359 | repE E106K, H120R, Q196R |
| 4 | R172C, H237R, K446E | | V266 | repE E106K, K107I |
| 6 | R337C | a466g | none | repE E101K, K107I |
| Final rescreen | I45V, P153S | - | I18, F251 | - |
| Final rescreen | V294A | - | A47, N111, I232, G359 | - |
| Final rescreen | E40K | - | F176 | GWS1B H464R (His-tag) |

**Supplementary Table 4.** Kinetic data for active variants (see Excel file).

**Supplementary Table 5.** GWS1B variant comparisons with *Nicotiana tabacum* Rubisco and *Rhodospirillum rubrum* Rubisco (see Excel file).

**Supplementary Table 6.** Plasmids and strains used in this study.

| Plasmid | Genes encoded | Source |
|---|---|---|
| pACYC$^{prk:nptII}$ | *prk* (WP_011242879.1), *nptII* (WP_000572405.1) | (1) |
| BAC-GWS1B | *Gallionellaceae rbcL* (ALP32085.1) | This work |
| pTrc-GWS1B | *Gallionellaceae rbcL* (ALP32085.1) | This work |
| **Strain** | **Genes encoded** | **Source** |
| NEB® 10-beta Competent *E. coli* (High Efficiency) | - | New England Biolabs |
| Dual8 *E. coli* (LMBP 13234) | MutaT7$^{C \rightarrow T}$, eMutaT7$^{A \rightarrow G}$ | (2) |

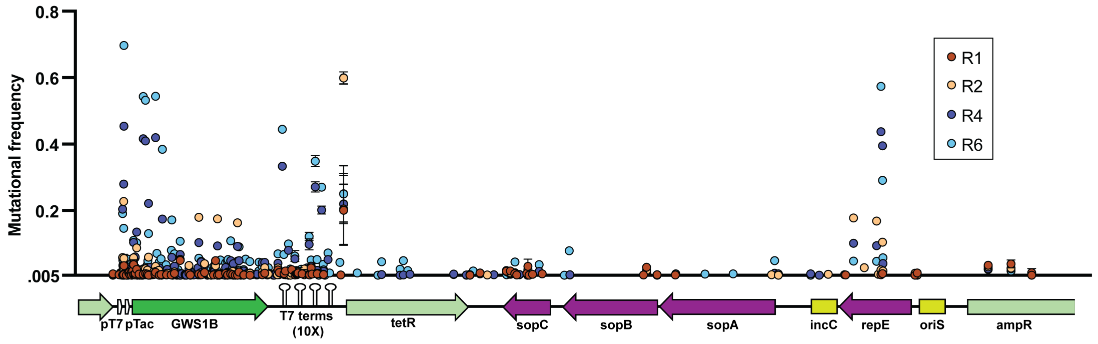


**Supplementary Figure 1.** Mutations identified in the coding regions of the BAC-GWS1B plasmid. See **Supplementary Table 2** for mutation details.

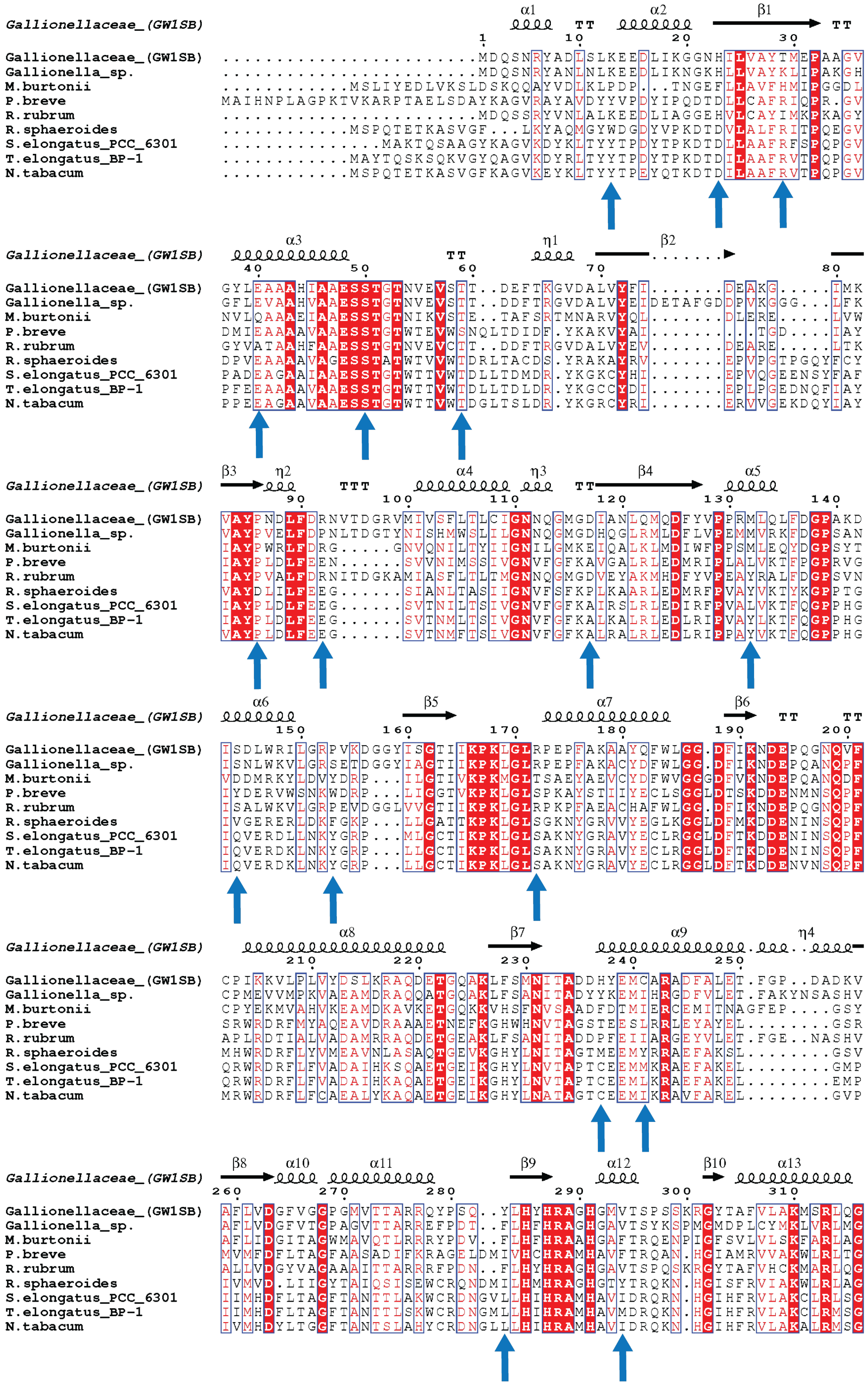
Gallionellaceae_(GW1SB) α1 TT α2 β1 TT
1 10 20 30
Gallionellaceae_(GW1SB) ........................MDQSNRYADLSLKEEDLIKGGNHILVAYTMEPAAGV
Gallionella_sp. ........................MDQSNRYANLNLKEEDLIKNGKHLLVAYKLIPAKGH
M.burtonii ............MSLIYEDLVKSLDSKQQAYVDLKLPDP...TNGEFLLAVFHMIPGGDL
P.breve MAIHNPLAGPKTVKARPTAELSDAYKAGVRAYAVDYYVPDYIPQDTDLLCAFRIQPR.GV
R.rubrum ........................MDQSSRYVNLALKEEDLIAGGEHVLCAYIMKPKAGY
R.sphaeroides ............MSPQTETKASVGF...LKYAQMGYWDGDYVPKDTDVLALFRITPQEGV
S.elongatus_PCC_6301 ..............MAKTQSAAGYKAGVKDYKLTYYTPDYTPKDTDLLAAFRFSPQPGV
T.elongatus_BP-1 ............MAYTQSKSQKVGYQAGVKDYRLTYYTPDYTPKDTDILAAFRVTPQPGV
N.tabacum ............MSPQTETKASVGFKAGVKEYKLTYYTPEYQTKDTDILAAFRVTPQPGV
Gallionellaceae_(GW1SB) α3 TT η1 β2
40 50 60 70 80
Gallionellaceae_(GW1SB) GYLEAAAHIAAESSTGTNVEVSTT..DEFTKGVDALVYFI.......DEAKG.....IMK
Gallionella_sp. GFLEVAAHVAAESSTGTNVEVSTT..DDFTRGVDALVYEIDETAFGDDPVKGGG...LFK
M.burtonii NVLQAAAEIAAESSTGTNIKVSTE..TAFSRTMNARVYQL.......DLERE.....LVW
P.breve DMIEAAAAVAAESSTGTWTEVWSNQLTDIDF.YKAKVYAI..........TGD....IAY
R.rubrum GYVATAAHFAAESSTGTNVEVCTT..DDFTRGVDALVYEV.......DEARE.....LTK
R.sphaeroides DPVEAAAAVAGESSTATWTVVWTDRLTACDS.YRAKAYRV.......EPVPGTPGQYFCY
S.elongatus_PCC_6301 PADEAGAAIAAESSTGTWTTVWTDLLTDMDR.YKGKCYHI.......EPVQGEENSYFAF
T.elongatus_BP-1 PFEEAAAAVAAESSTGTWTTVWTDLLTDLDR.YKGCCYDI.......EPLPGEDNQFIAY
N.tabacum PPEEAGAAVAAESSTGTWTTVWTDGLTSLDR.YKGRCYRI.......ERVVGEKDQYIAY
Gallionellaceae_(GW1SB) β3 η2 TTT α4 η3 TT β4 α5
90 100 110 120 130 140
Gallionellaceae_(GW1SB) VAYPNDLFDRNVTDGRVMIVSFLTLCIGNNQGMGDIANLQMQDFYVPPRMLQLFDGPAKD
Gallionella_sp. VAYPVELFDPNLTDGTYNISHMWSLILGNNQGMGDHQGLRMLDFLVPEMMVRKFDGPSAN
M.burtonii IAYPWRLFDRG.....GNVQNILTYIIGNILGMKEIQALKLMDIWFPPSMLEQYDGPSYT
P.breve IAYPLDLFEEN......SVNIMSSIVGNVFGFKAVGALRLEDMRIPLALVKTFPGPRVG
R.rubrum IAYPVALFDRNITDGKAMIASFLTLTMGNNQGMGDVEYAKMHDFYVPEAYRALFDGPSVN
R.sphaeroides VAYDLILFEEG......SIANLTASIIGNVFSFKPLKAARLEDMRFPVAYVKTYKGPPTG
S.elongatus_PCC_6301 IAYPLDLFEEG......SVTNILTSIVGNVFGFKALRSLRLEDIRFPVALVKTFQGPPHG
T.elongatus_BP-1 IAYPLDLFEEG......SVTNMLTSIVGNVFGFKALKALRLEDLRIPVAYLKTFQGPPHG
N.tabacum VAYPLDLFEEG......SVTNMFTSIVGNVFGFKALRALRLEDLRIPPAYVKTFQGPPHG
Gallionellaceae_(GW1SB) α6 β5 α7 β6 TT TT
150 160 170 180 190 200
Gallionellaceae_(GW1SB) ISDLWRILGRPVKDGGYISGTIIKPKLGLRPEPFAKAAYQFWLGG.DFIKNDEPQGNQVF
Gallionella_sp. ISNLWKVLGRSETDGGYIAGTIIKPKLGLRPEPFAKACYDFWLGG.DFIKNDEPQANQPF
M.burtonii VDDMRKYLDVYDRP...ILGTIVKPKMGLTSAEYAEVCYDFWVGGGDFVKNDEPQANQDF
P.breve IYDERVWSNKWDRP...LIGGTVKPKLGLSPKAYSTIIYECLSGGLDTSKDDENMNSQPF
R.rubrum ISALWKVLGRPEVDGGLVVGTIIKPKLGLRPKPFAEACHAFWLGG.DFIKNDEPQGNQPF
R.sphaeroides IVGERERLDKFGKP...LLGATTKPKLGLSGKNYGRVVYEGLKGGLDFMKDDENINSQPF
S.elongatus_PCC_6301 IQVERDLLNKYGRP...MLGCTIKPKLGLSAKNYGRAVYECLRGGLDFTKDDENINSQPF
T.elongatus_BP-1 IQVERDKLNKYGRP...LLGCTIKPKLGLSAKNYGRAVYECLRGGLDFTKDDENINSQPF
N.tabacum IQVERDKLNKYGRP...LLGCTIKPKLGLSAKNYGRAVYECLRGGLDFTKDDENVNSQPF
Gallionellaceae_(GW1SB) α8 β7 α9 η4
210 220 230 240 250
Gallionellaceae_(GW1SB) CPIKKVLPLVYDSLKRAQDETGQAKLFSMNITADDHYEMCARADFALET.FGP..DADKV
Gallionella_sp. CPMEVVMPKVAEAMDRAQQATGQAKLFSANITADYYKEMIHRGDFVLET.FAKYNSASHV
M.burtonii CPYEKMVAHVKEAMDKAVKETGQKKVHSFNVSAADFDTMIERCEMITNAGFEP....GSY
P.breve SRWRDRFMYAQEAVDRAAAETNEFKGHWHNVTAGSTEESLRRLEYAYEL........GSR
R.rubrum APLRDTIALVADAMRRAQDETGEAKLFSANITADDPFEIIARGEYVLET.FGE..NASHV
R.sphaeroides MHWRDRFLYVMEAVNLASAQTGEVKGHYLNITAGTMEEMYRRAEFAKSL........GSV
S.elongatus_PCC_6301 QRWRDRFLFVADAIHKSQAETGEIKGHYLNVTAPTCEEMMKRAEFAKEL........GMP
T.elongatus_BP-1 QRWRDRFLFVADAIHKAQAETGEIKGHYLNVTAPTCEEMLKRAEFAKEL........EMP
N.tabacum MRWRDRFLFCAEALYKAQAETGEIKGHYLNATAGTCEEMIKRAVFAREL........GVP
Gallionellaceae_(GW1SB) β8 α10 α11 β9 α12 β10 α13
260 270 280 290 300 310
Gallionellaceae_(GW1SB) AFLVDGFVGGPGMVTTARRQYPSQ..YLHYHRAGHGMVTSPSSKRGYTAFVLAKMSRLQG
Gallionella_sp. AFLVDGFVTGPAGVTTARRFEPDT..FLHFHRAHGGAVTSYKSPMGMDPLCYMKLVRLMG
M.burtonii AFLIDGITAGWMAVQTLRRRYPDV..FLHFHRAAHGAFTRQENPIGFSVLVLSKFARLAG
P.breve MVMFDFLTAGFAASADIFKRAGELDMIVHCHRAMHAVFTRQAN.HGIAMRVVAKWLRLTG
R.rubrum ALLVDGYVAGAAAITTARRRFPDN..FLHYHRAGHGAVTSPQSKRGYTAFVHCKMARLQG
R.sphaeroides IVMVD.LIIGYTAIQSISEWCRQNDMILHMHRAGHGTYTRQKN.HGISFRVIAKWLRLAG
S.elongatus_PCC_6301 IIMHDFLTAGFTANTTLAKWCRDNGVLLHIHRAMHAVIDRQRN.HGIHFRVLAKCLRLSG
T.elongatus_BP-1 IIMHDFLTAGFTANTTLSKWCRDNGMLLHIHRAMHAVMDRQKN.HGIHFRVLAKCLRMSG
N.tabacum IVMHDYLTGGFTANTSLAHYCRDNGLLLHIHRAMHAVIDRQKN.HGIHFRVLAKALRMSG

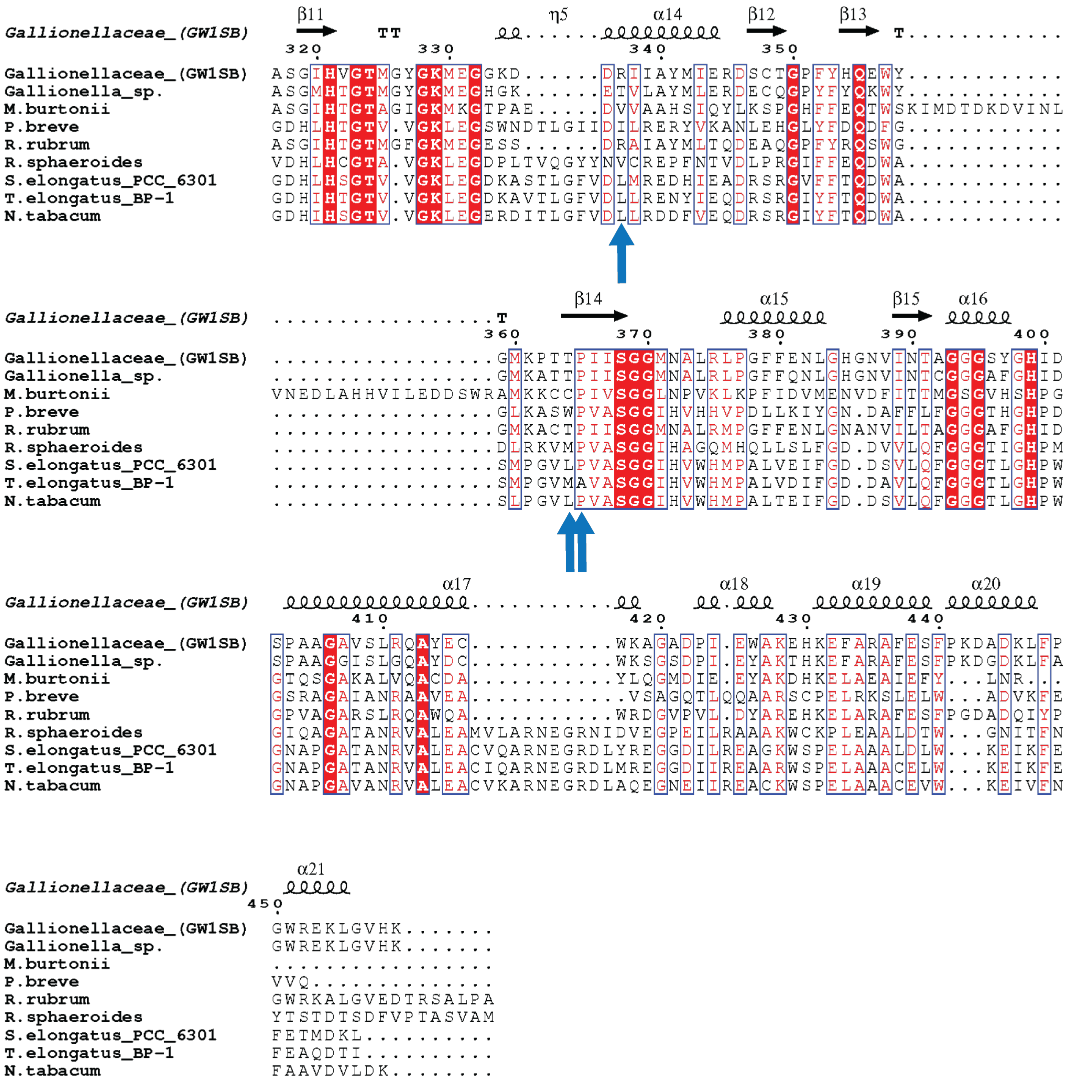


**Supplementary Figure 2.** Protein sequence alignment of GWS1B with diverse Rubisco isoforms created using ESPript 3.0 (3). GWS1B secondary structure is displayed above alignment. Conserved residues are highlighted in red. Substitutions identified via next-generation sequencing are indicated with blue arrows.

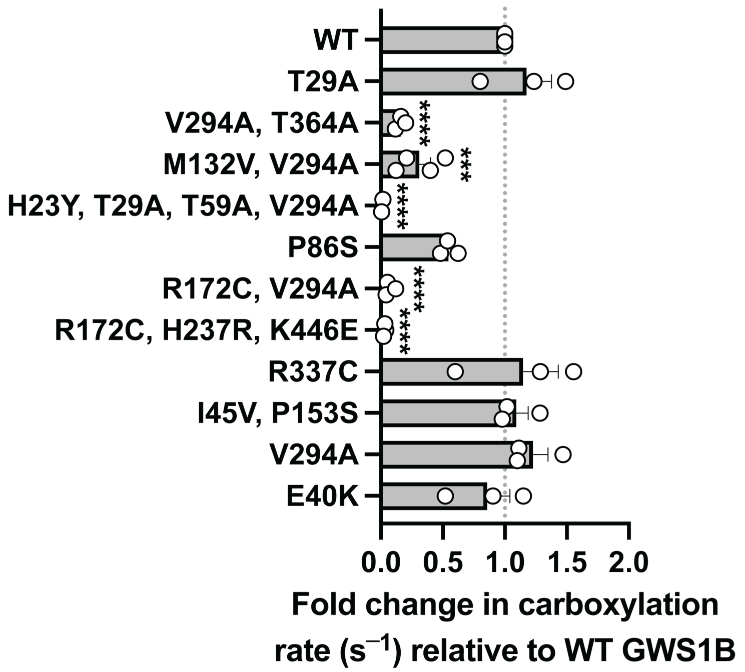


**Supplementary Figure 3.** Spectrophotometric determination of carboxylation rate at 21% $O_2$ relative to wild-type GWS1B for all variants. The data are the means and standard deviations of three replicates, with significance shown relative to wild-type GWS1B analyzed by one-way ANOVA. ***, $p \leq .0005$; ****, $p \leq .00005$.

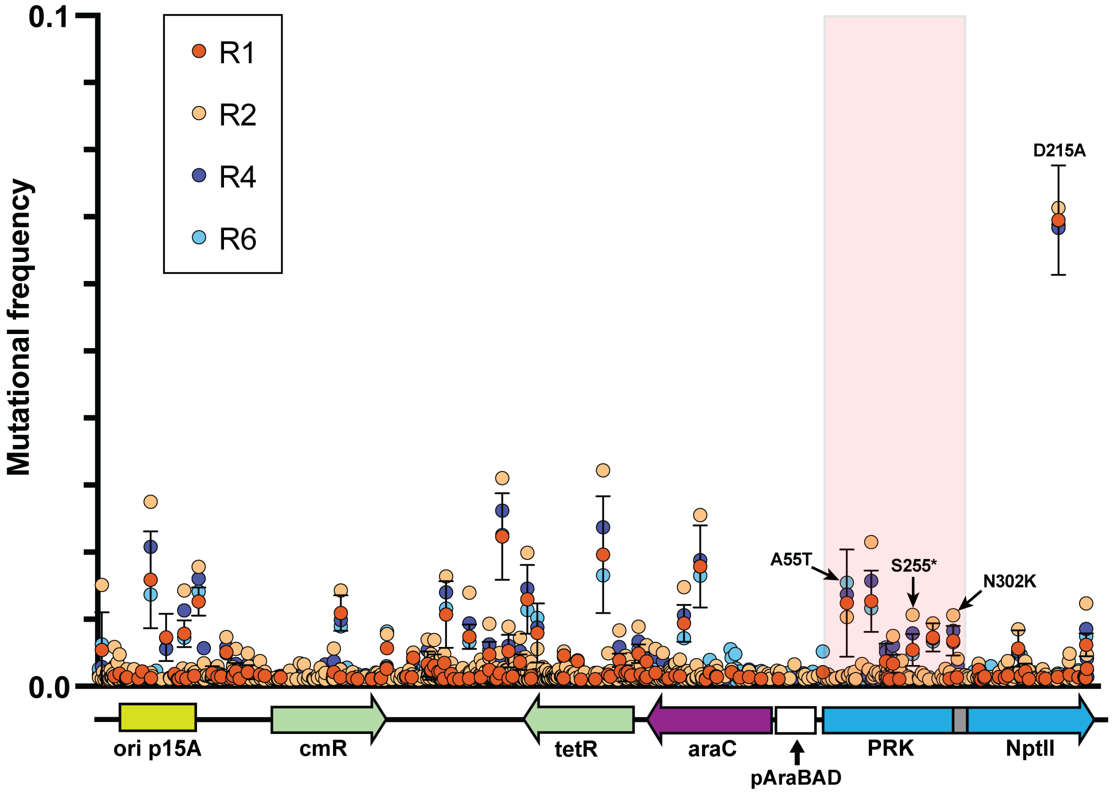


**Supplementary Figure 4.** Mutations identified in all coding regions of pACYC$^{prk:nptII}$. PRK is shaded red. High frequency, non-synonymous mutations in PRK-NPTII are labeled.

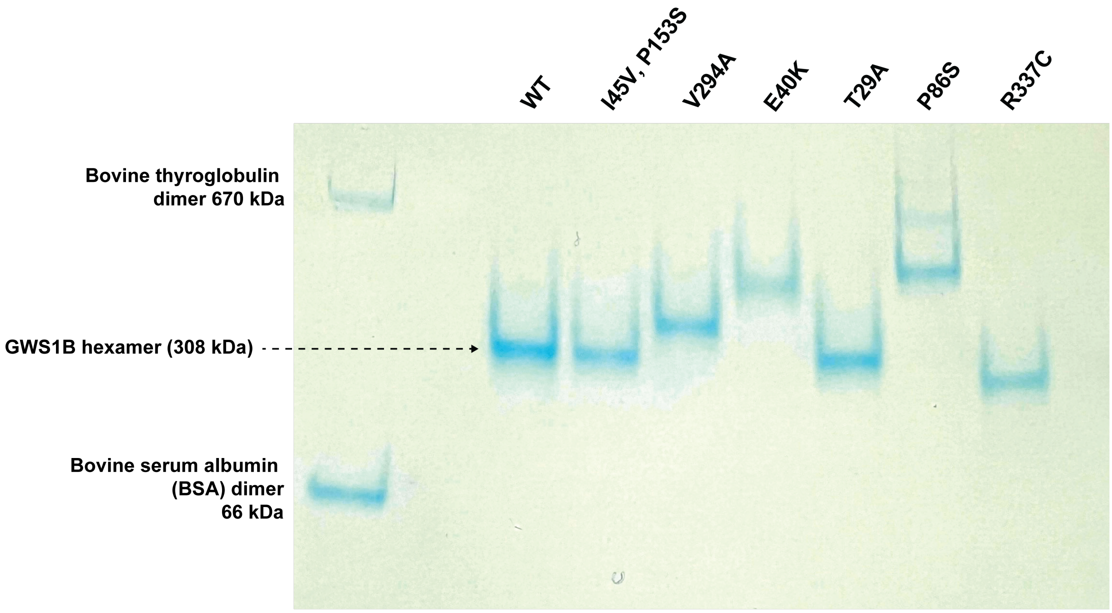


**Supplementary Figure 5.** Native PAGE of GWS1B variants. The presence of faint, higher molecular weight bands likely indicates higher-order oligomers or aggregates. Changes in extent of migration are most likely due to charge changes or small radii changes.

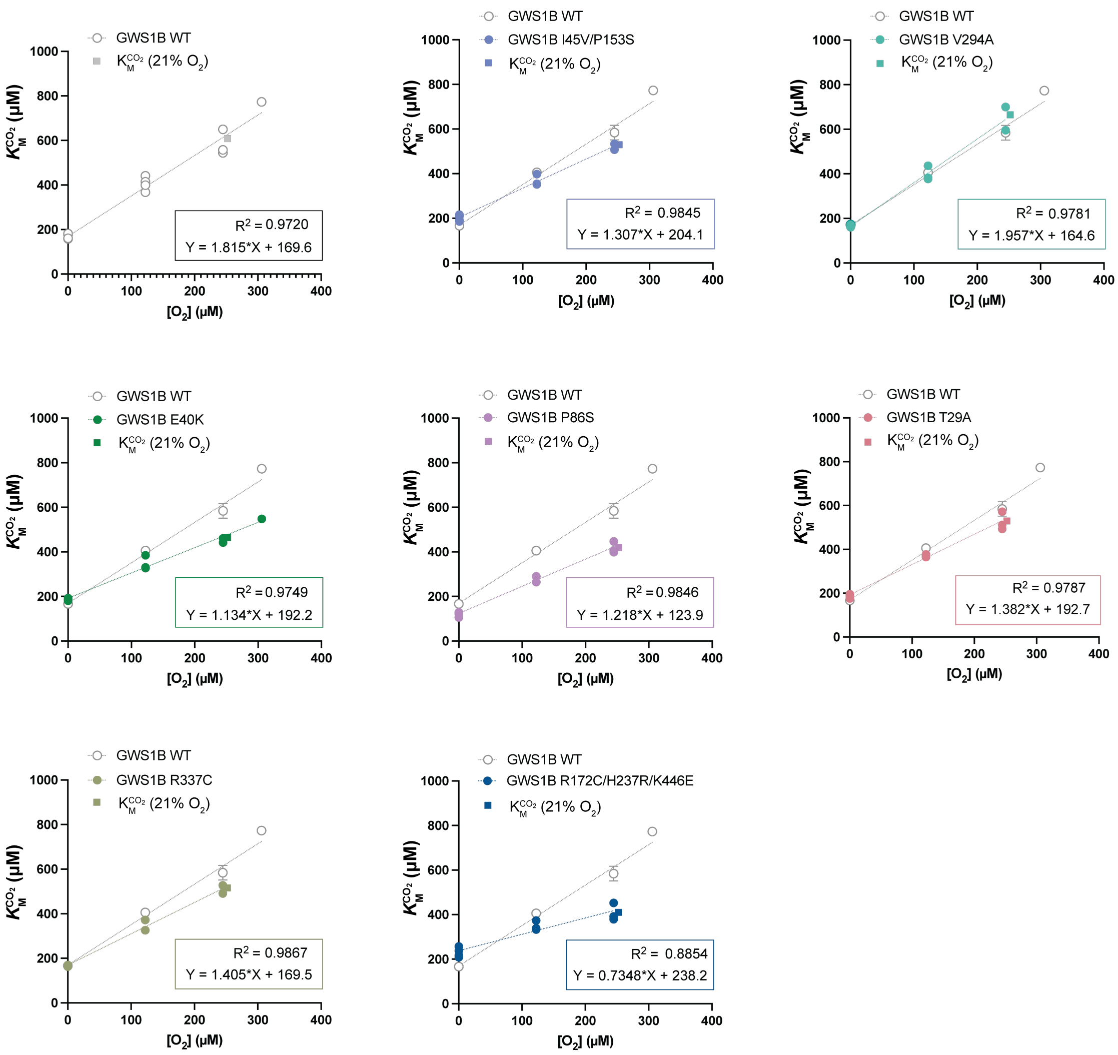


**Supplementary Figure 6.** Determination of $K_M^{O_2}$ and $K_M^{CO_2}$ $(21\%\ O_2)$. $K_M^{CO_2}$ of wild-type and variant GWS1B enzymes were measured over a range of $O_2$ concentrations. $K_M^{O_2}$ and $K_M^{CO_2}$ $(21\%\ O_2)$ were calculated from the equation $K_{M\ app}^{CO_2} = K_M^{CO_2}\ (1 + \frac{[O_2]}{K_M^{O_2}})$, or $y = K_M^{CO_2}\ (1 + \frac{x}{K_M^{O_2}})$, using 252 µM as 21% [$O_2$] in air-saturated $H_2O$ (as assays performed at 983 mbar in Canberra, Australia). Inset shows the $R^2$ value of linear fit equation with the extrapolated $K_M^{CO_2}$ $(21\%\ O_2)$ value indicated by a rectangle.

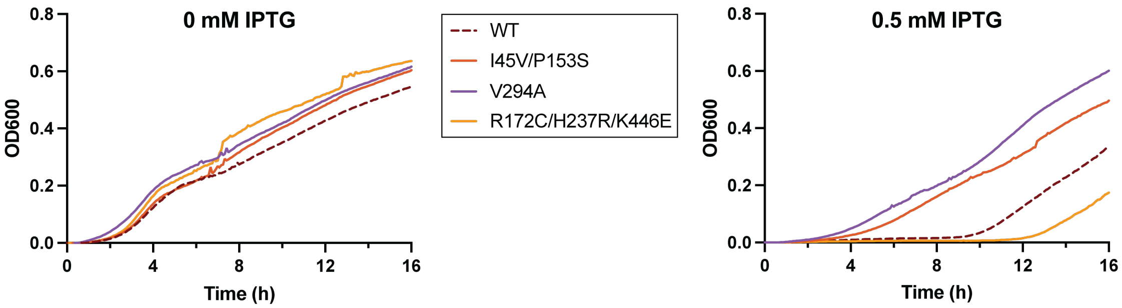


**Supplementary Figure 7.** Growth curves ($OD_{600}$) measured over 16 h at 37 °C of wild-type, I45V/P153S, V294A, or R172C/H237R/K446E pTrc-GWS1B transformed *E. coli* under GWS1B Rubisco non-inducing (0 mM IPTG) or inducing (0.5 mM IPTG) conditions.
